# Genes relocated between *Drosophila chromosome* arms evolve under relaxed selective constraints relative to non-relocated genes

**DOI:** 10.1101/178582

**Authors:** Margaret L. I. Hart, Ban L. Vu, Quinten Bolden, Keith T. Chen, Casey L. Oakes, Lejla Zoronjic, Richard P. Meisel

## Abstract

Gene duplication creates a second copy of a gene either in tandem to the ancestral locus or dispersed to another chromosomal location. When the ancestral copy of a dispersed duplicate is lost from the genome, it creates the appearance that the gene was “relocated” from the ancestral locus to the derived location. Gene relocations may be as common as canonical dispersed duplications in which both the ancestral and derived copies are retained. Relocated genes appear to be under more selective constraints than the derived copies of canonical duplications, and they are possibly as conserved as single-copy non-relocated genes. To test this hypothesis, we combined comparative genomics, population genetics, gene expression, and functional analyses to assess the selection pressures acting on relocated, duplicated, and non-relocated single-copy genes in *Drosophila* genomes. We find that relocated genes evolve faster than single-copy non-relocated genes, and there is no evidence that this faster evolution is driven by positive selection. In addition, relocated genes are less essential for viability and male fertility than single-copy non-relocated genes, suggesting that relocated genes evolve fast because of relaxed selective constraints. However, relocated genes evolve slower than the derived copies of canonical dispersed duplicated genes. We therefore conclude that relocated genes are under more selective constraints than canonical duplicates, but are not as conserved as single-copy non-relocated genes.

## Introduction

Duplicated genes are important contributors to molecular evolution (Ohno, 1970; Conant and Wolfe, 2008; Dittmar and Liberles, 2010; Innan and Kondrashov, 2010). A gene duplication event creates a second (derived) copy of a gene via one of many molecular mechanisms, including non-allelic recombination and reverse transcription of mRNA (Zhang, 2003; Kaessmann *et al.*, 2009; Marques-Bonet *et al.*, 2009). The derived copy can acquire novel functions and/or the ancestral and derived loci can each evolve a subset of functions present prior to duplication (Spofford, 1969; Hughes, 1994; Force *et al.*, 1999; Lynch and Force, 2000). When functions are partitioned between the paralogous copies, gene duplication can resolve pleiotropic conflicts present in the single-copy ancestor (Hittinger and Carroll, 2007; Des Marais and Rausher, 2008; Connallon and Clark, 2011; Gallach and Betrán, 2011; Abascal *et al.*, 2013; VanKuren and Long, 2018).

**Figure 1:**
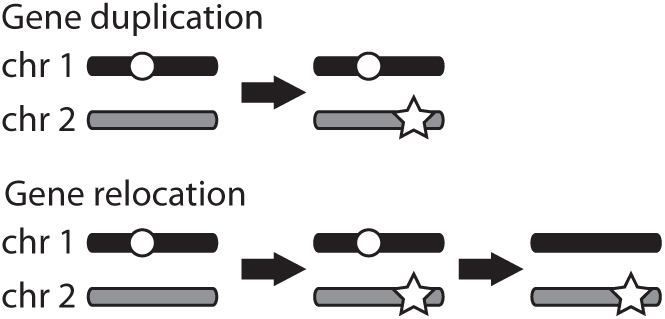
Gene duplication and relocation. In the ancestral arrangement, a gene (white circle) is located on chromosome 1. After gene duplication, the derived copy (star) is located on chromosome 2. In the case of gene relocation, the copy at the ancestral locus is subsequently lost.

Gene duplication can give rise to a derived copy located in tandem to the ancestral copy or dispersed to another genomic location. The ancestral copy of a dispersed duplicate can be lost from the genome, creating the appearance that the gene was “relocated” to the derived locus (**Fig 1**). Comparative genomic analyses in animals and plants have revealed that gene relocation occurs frequently, and relocated genes may be as common as canonical dispersed duplications in which the ancestral copy is retained (Bhutkar *et al.*, 2007; Meisel *et al.*, 2009; Wicker *et al.*, 2010; Han and Hahn, 2012; Ciomborowska *et al.*, 2013). Furthermore, gene relocation can promote reproductive isolation between species because some F_2_ hybrids lack the relocated gene (Masly *et al.*, 2006; Bikard *et al.*, 2009; Moyle *et al.*, 2010).

Despite the prevalence and evolutionary importance of gene relocation, the selection pressures acting on relocated genes have received considerably less attention than the evolutionary dynamics of canonical duplicated genes. The analyses that have been performed identified some important differences between relocated genes and canonical dispersed duplicates. For example, derived copies of duplicated genes in animal genomes tend to be narrowly expressed in reproductive tissues (Vinckenbosch *et al.*, 2006; Meisel *et al.*, 2009, 2010; Baker *et al.*, 2012; Kondo *et al.*, 2017). In contrast, *Drosophila* and human relocated genes tend to be broadly expressed across many tissues (Meisel *et al.*, 2009; Ciomborowska *et al.*, 2013). In addition, the derived copies of dispersed duplicates tend to experience positive selection or relaxed constraints (Kondrashov *et al.*, 2002; Conant and Wagner, 2003; Han *et al.*, 2009; Han and Hahn, 2012), while mammalian relocated genes appear to evolve under strong purifying selection (Ciomborowska *et al.*, 2013). It has been hypothesized that positive selection on the derived copies of duplicated genes fixes mutations that improve testis-specific functions once pleiotropic constraints are relaxed by duplication (Betrán and Long, 2003; Torgerson and Singh, 2004; Betrán *et al.*, 2006; Rosso *et al.*, 2008; Meisel *et al.*, 2010; Quezada-Diaz *et al.*, 2010; Tracy *et al.*, 2010; VanKuren and Long, 2018). Gene relocation is unlikely to resolve pleiotropic conflicts because a second copy of the gene is not retained. To improve our understanding of the evolutionary dynamics of relocated genes, we combined population genetic, functional genomic, and experimental approaches to characterize the selection pressures acting on *Drosophila* relocated genes.

## Materials and Methods

### Identifying duplicated and relocated genes

*Drosophila* genomes have six chromosome arms, known as Muller elements A–F (Muller, 1940; Schaeffer *et al.*, 2008). We analyzed previously annotated inter-chromosome-arm duplicated and relocated genes that occurred along the lineages leading to *Drosophila melanogaster* and *Drosophila pseudoobscura* (Hahn *et al.*, 2007; Meisel *et al.*, 2009), ignoring duplication and relocation events involving the minute element F. The lineage-specific duplicates were identified by examining phylogenetic reconstructions of gene families from the *D. melanogaster, D. pseudoobscura, Drosophila willistoni, Drosophila virilis*, and *Drosophila grimshawi* genomes. We selected gene families in which the phylogenetic reconstruction included a duplication event along the lineage leading to *D. melanogaster* or *D. pseudoobscura* after the divergence with all other lineages. From this group, we then curated a list of duplications in which one copy was on a different chromosome arm than the homologous genes across all species. The ancestral copy of a duplicated gene in one species’ genome was inferred to be the copy found on the same chromosome arm as in the other four species, and the derived copy is the one on a different chromosome arm. Relocated genes were identified as present in a single copy in *D. melanogaster* or *D. pseudoobscura*, with single-copy orthologs on a different chromosome arm in the other four species. As a control, we also analyzed singlecopy non-relocated genes that are retained as 1:1:1:1:1 orthologs on the same chromosome arm across all five species (Meisel *et al.*, 2009). We excluded genes on element F from our control set.

### Sequence divergence, polymorphism, and selection

We obtained estimates of polymorphism and divergence for relocated genes, non-relocated single-copy genes, and the ancestral and derived copies of inter-chromosome-arm duplicated genes in the *D. melanogaster* genome from published datasets. Two data sets were used to calculate divergence between *D. melanogaster* and *Drosophila simulans* orthologs. All of the duplications and relocations in our data set happened before the divergence of the *D. melanogaster* and *D. simulans* lineages, so that our estimates of divergence are specific to either the ancestral or derived copy. First, we obtained estimates of nucleotide sequence divergence along the *D. melanogaster* lineage after the split with *D. simulans* for all 1:1 orthologous genes between these two closely related species (Hu *et al.*, 2013). In the results presented here, we analyzed substitutions per site for 0-fold and 4-fold degenerate sites within protein coding regions. Second, we obtained estimates of the ratio of non-synonymous to synonymous substitutions per site (*d*_*N*_/*d*_*S*_) from a published analysis comparing *D. melanogaster* and *D. simulans* genes (Stanley and Kulathinal, 2016).

We obtained the amount non-synonymous (*P*_*N*_) and synonymous (*P*_*S*_) polymorphic sites within *D. melanogaster* genes from the *Drosophila* Genetic Reference Panel (DGRP; Mackay *et al.*, 2012; Ràmia *et al.*, 2012). We only included polymorphic sites with a minor allele frequency *>*5% to minimize the inclusion of segregating deleterious alleles (Fay *et al.*, 2001). We also obtained the number of non-synonymous (*D*_*N*_) and synonymous (*D*_*S*_) substitutions between *D. melanogaster* and *D. simulans* from the DGRP data. We analyzed the polymorphism and divergence data for single-copy non-relocated genes, the ancestral and derived copies of inter-chromosome-arm duplicates, and relocated genes within the framework of McDonald and Kreitman (1991). First, we used a *?*^2^ test of independence to identify genes with an excess or deficiency of non-synonymous substitutions. We assigned genes as evolving under positive selection if they have a significant excess of non-synonymous substitutions, and we assigned genes as evolving under strong negative selection if they have a significant deficiency of non-synonymous substitutions. Second, we calculated 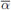the fraction of nonsynonymous substitutions fixed by selection (Smith and Eyre-Walker, 2002), for each group of genes:

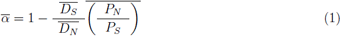

We calculated 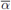separately for single-copy non-relocated genes, ancestral copies of inter-chromosome-arm duplicates, derived copies, and relocated genes. We performed 1,000 boot-strapped replicate analyses to calculate a confidence interval (CI) for each 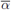estimate.

### Gene expression profiles

We analyzed available microarray data to assess the expression across adult tissues of *D. melanogaster* relocated genes, duplicated genes, and single-copy genes. Expression measurements were taken from FlyAtlas, which includes 11 non-redundant adult non-sex-specific tissue samples (brain, crop, midgut, hindgut, Malpighian tubule, thoracicoabdominal ganglion, salivary gland, fat body, eye, heart, and trachea), two male-specific organs (testis and accessory gland), and two female-specific organs (ovary and spermatheca) (Chintapalli *et al.*, 2007). Expression levels for spermatheca were averaged between mated and unmated females (Meisel, 2009). We used *τ* as a measure of expression breadth for each gene:

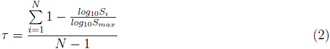

where *N* is the number of tissues (15), *S*_*i*_ is the expression level in tissue *i*, and *S*_*max*_ is the maximum expression of that gene across all tissues (Yanai *et al.*, 2005; Larracuente *et al.*, 2008). All *S*_*i*_1 were set to 1 for this analysis. Values of *τ* range from 0 to 1, with higher values corresponding to more tissue-specific expression.

We also analyzed microarray data from *D. melanogaster* testis (Chintapalli *et al.*, 2007) and RNA-seq data from *D. pseudoobscura* testis (Meisel *et al.*, 2010) to infer the expression levels of relocated, duplicated, and single-copy genes. Finally, we analyzed sex-specific microarray data from *D. melanogaster* and *D. pseudoobscura* heads and whole flies to calculate “sex-biased” expression (Meisel *et al.*, 2012), i.e., the relative expression of genes in males and females 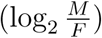

### Viability and fertility effects of knockdown

To assess if relocated and non-relocated single-copy *D. melanogaster* genes are essential for viability and male fertility, we used Gal4-UAS inducible RNA interference (RNAi) to knock down the expression of relocated and single-copy non-relocated genes. Flies carrying an inducible construct containing a hairpin sequence that silences the expression of a target gene via RNAi (UAS-RNAi) were obtained from the Vienna Drosophila Resource Center (VDRC; Dietzl *et al.*, 2007). Knockdown was performed using two different sets of RNAi lines. The first set, known as “GD” lines, were produced by random integration into the *D. melanogaster* genome of a P-element construct carrying a pUAST vector with 10 copies of the UAS and a 300–400bp inverted repeat targeting the gene of interest. The second set, known as “KK” lines, also carry 10 copies of UAS and a long inverted repeat, but they were inserted into specific sites in the genome using ϕC31 targeted integration (Groth *et al.*, 2004; Bateman *et al.*, 2006). Expression of the RNAi construct in some of the KK lines can be lethal because of mis-expression of the developmental gene *tiptop* (Green *et al.*, 2014; Vissers *et al.*, 2016), which can lead to false inference about the essentiality of duplicated genes (Kondo *et al.*, 2017). We therefore performed analyses of our results from the GD and KK lines separately to assess the extent to which our results could be attributed to systemic effects of KK lines.

To assay the effect of knockdown on viability, individual males carrying a UAS-RNAi transgene were crossed to individual females carrying a Gal4 driver construct that is ubiquitously expressed under the *tubulin 1 α* promoter (*P{tubP-Gal4}*). *P{tubP-Gal4}* is expressed in many tissues and throughout development (Lee and Luo, 1999), which causes constitutive knockdown of the target gene when combined in the same genotype with a UAS-RNAi construct. In addition, *P{tubP-Gal4}* is balanced over the TM3 chromosome, which carries the dominant *Stubble* (*Sb*) allele, allowing us to differentiate between knockdown and non-knockdown (control) siblings within each cross. The females were allowed to lay eggs for 3 days following mating on cornmeal media, and then all progeny that emerged were scored for their sex and bristle phenotype (stubble or wild-type). We assessed the viability of the knockdown flies by comparing the counts of knockdown progeny with their control siblings. We also performed control crosses in which the UAS-RNAi male is replaced with a male from the progenitor stock from which the RNAi lines were derived—GD lines were created by transforming *w* ^1118^ flies (VDRC line 60000), and KK lines were created by transforming *y,w* ^1118^;*P{*attP,*y* ^+^,*w* ^3^’*}* flies (VDRC line 60100). These control males do not carry a UAS-RNAi construct.

We used linear models to assess the effect of RNAi knockdown on viability. For each gene, we modeled the number of progeny recovered (*N*_*ijk*_) with the phenotype associated with either knockdown (wild-type bristles) or non-knockdown (stubble bristles) from crosses involving a fly either carrying the UAS-RNAi construct or from the progenitor non-RNAi strain:

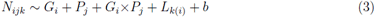

where *G*_*i*_ is a fixed effect indicating if the line carried a UAS-RNAi construct targeting the gene or if it was a control line; *P*_*j*_ is a fixed effect indicating the phenotype of the progeny (either knockdown or stubble control); *L*_*k(i*_) is a fixed effect (nested within *G*_*i*_) indicating the UAS-RNAi construct used to knock down the target gene; and *b* is a random effect indicating the replicate block in which the viability assay was performed. If only one UASRNAi construct was used to knock down the target gene, then *L*_*k(i)*_ was excluded from equation 3. The effect of knockdown on viability was estimated as the effect of the *G*_*i*_×*P*_*j*_ interaction for crosses in which the gene is knocked down and the progeny have wild-type bristles. If the *G*_*i*_×*P*_*j*_ interaction has a significant effect on the number of progeny recovered from the cross (*N*_*ijk*_), then there is an effect of knockdown on viability. To test for significance of the interaction term, we used a drop in deviance test to compare the fit of the full model with a model excluding the interaction term.

To assess the effects of RNAi knockdown on male fertility, we crossed UAS-RNAi males to females carrying a Gal4 driver construct that is constitutively expressed under the *bag of marbles* (*bam*) promoter (*P{bam-Gal4-VP16}*) to create male progeny in which the target gene is knocked down the germline (Sartain *et al.*, 2011). The *bam* promoter drives expression in germ cells after differentiation from the stem cells (Chen and McKearin, 2003). We assessed the fertility of the knockdown male progeny by crossing individual 5 day old males to single 4–6 day old Orergon R (OreR) or Canton S (CanS) virgin females, and we observed all matings to ensure that copulation occurred. We only considered the results of matings in which we observed copulation between the male and the CanS/OreR female to ensure that the fertility assay was not confounded by behavioral effects that interfere with mating success. The females were allowed to lay eggs for 2 days after mating on cornmeal media, they were then transferred to a new vial for 2 additional days of egg laying, and the total number of adult progeny that emerged in both vials were added together as a measure of the fertility of the knockdown male. As a control for each batch, we assessed the fertility of males that were created by crossing *bam-Gal4* females with males from the progenitor strains that do not carry the UAS-RNAi constructs.

For each gene, we modeled the number of progeny recovered (*N*_*ijk*_) from matings involving either a male carrying the UAS-RNAi construct or a control male carrying a chromosome from the progenitor line:

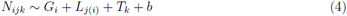

where *G*_*i*_ is a fixed effect indicating if the male carried a UAS-RNAi construct targeting the gene or if it was a control line; *L*_*j(i)*_ is a fixed effect (nested within *G*_*i*_) indicating the UAS-RNAi construct used to knock down the target gene; *T*_*k*_ is a fixed effect indicating the genotype of the female used to assess fertility (either CanS or OreR); and *b* is a random effect indicating the replicate block in which the fertility assay was performed. If only one UAS-RNAi construct was used to knock down the target gene, then *L*_*j*_(*i*) was excluded from equation 4. The effect of knockdown on fertility is quantified by *G*_*i*_. If *G*_*i*_ has a significant effect on the number of progeny (*N*_*ijk*_), then there is an effect of germline knockdown on male fertility. To test for a significant effect of male genotype, we used a drop in deviance test to compare the fit of the full model with a model excluding the male genotype (*G*_*i*_). For some of the genes, only one UAS-RNAi line and one female genotype were used in a single block, and we therefore could not use the drop in deviance to assess the effect of knockdown on fertility. In those cases, we assessed the effect of the male genotype using a single factor ANOVA (equivalent to a Student’s T-test): *N*_*i*_ ∼ *G*_*i*_. All analyses were performed in the R statistical programming environment (R Core Team, 2015).

## Data availability

All divergence data, gene expression data, and results from RNAi experiments are available as supplemental files. File S1 contains a description of all supplemental data.

## Results

### Relocated genes evolve fast because of relaxed selective constraints

We tested if the protein coding sequences of genes that were relocated to other chromosome arms along the *D. melanogaster* lineage evolve at different rates than inter-chromosome-arm duplicated genes or single-copy non-relocated genes. The ancestral copies of duplicated genes, derived copies, and relocated genes all evolve faster at 0-fold degenerate (amino acid changing) sites than single-copy non-relocated genes (**Fig 2A**). Accelerated amino acid sequence evolution can be driven by positive selection, relaxed constraints, or higher mutation rates. The derived copies of duplicated genes evolve faster than single-copy genes at 4-fold degenerate (silent) sites (**Fig 2B**), suggesting that higher mutation rates could explain the faster evolution of derived copies at 0-fold degenerate sites. However, *d*_*N*_ /*d*_*s*_ is significantly elevated in the ancestral copies, derived copies, and relocated genes relative to single-copy non-relocated genes (**Fig 2C**). We therefore conclude that mutational bias cannot entirely explain the faster amino acid sequence evolution of relocated genes.

Other analyses have found that the derived copies of duplicated genes evolve faster and experience more positive selection than the ancestral copies (Kondrashov *et al.*, 2002; Conant and Wagner, 2003; Han *et al.*, 2009; Han and Hahn, 2012). We fail to detect significant differences between ancestral and derived copies of inter-chromosome-arm duplicated genes in divergence at 0-fold degenerate sites (*P* =0.373), divergence at 4-fold degenerate sites (*P* =0.553), or *d*_*N*_ /*d*_*S*_ (*P* =0.208; all *P* values from Mann-Whitney *U* tests). However, we have small sample sizes of ancestral and derived duplicates (14–30 depending on the divergence estimate), which likely limits our power to detect significant differences in evolutionary rates. There are substantially more relocated genes with divergence estimates (34–60), and we detect significantly elevated *d*_*N*_ /*d*_*S*_ in the derived copies of duplicated genes relative to relocated genes (*P* =7.5*×*10^−5^ in a Mann-Whitney *U* test). Our results demonstrate that the protein coding sequences of relocated genes evolve faster than single-copy non-relocated genes, and the derived copies of duplicated genes evolve faster than relocated genes.

**Figure 2:**
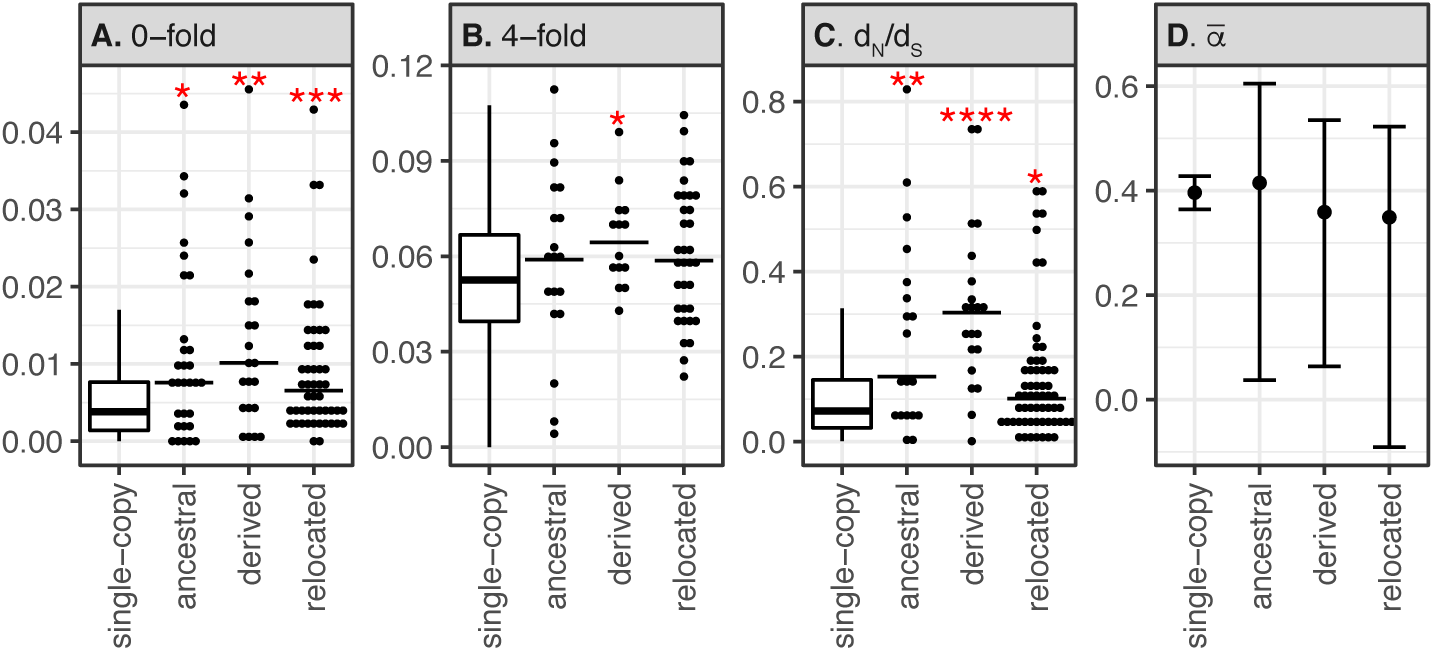
Divergence and polymorphism-divergence statistics for single-copy non-relocated genes, the ancestral and derived copies of inter-chromosome-arm duplicated genes, and relocated genes are plotted. Divergence estimates are between *D. melanogaster* and *D. simulans* at (A) 0-fold degenerate sites, (B) 4-fold degenerate sites, and (C) *d*_*N*_ /*d*_*S*_. The distribution of divergence values for single-copy genes is represented by a boxplot, while individual divergence values are shown for each of the other genes as a point (with the median indicated by a horizontal line). Significant differences in divergence when comparing single-copy genes with either ancestral copies, derived copies, or relocated genes are shown by red asterisks (^*^*P >* 0.05, ^**^*P >* 0.005, ^***^*P >* 0.0005, and ^****^*P >* 0.00005 in a Mann-Whitney *U* test). (D) Point estimates of *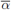* are plotted along with the 95% CI.

To distinguish between relaxed selective constraints (decreased purifying selection) and increased adaptive substitutions (positive selection) driving the rapid evolution of relocated genes, we analyzed polymorphism and divergence data. If accelerated evolutionary diver-gence is driven by positive selection, we expect the ratio of non-synonymous to synonymous substitutions to be greater than non-synonymous to synonymous polymorphisms (McDonald and Kreitman, 1991). Only a handful of duplicated and relocated genes have an excess of non-synonymous substitutions (**Table 1**). In addition, the proportion of ancestral copies, derived copies, or relocated genes with evidence for positive selection is not greater than the proportion of single-copy non-relocated genes with evidence for positive selection (**Table 1**). Furthermore, the fraction of non-synonymous substitutions fixed by selection (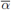) in relocated genes falls below the 95% CI for single-copy non-relocated genes (**Fig 2D**). There is also not a significant difference in 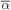 between relocated genes, ancestral copies of duplicated genes, or derived copies (**Fig 2D**). In summary, our results provide no evidence that relocated genes experience a disproportionate amount of positive selection, and we conclude that the accelerated evolution of relocated genes is driven by relaxed selective constraints.

**Table 1:** Counts of *D. melanogaster* single-copy non-relocated genes, ancestral copies of duplicated genes, derived copies, and relocated genes with no evidence for strong selection, evidence for positive selection, and evidence for negative selection.

|  | no selection | positive selection | negative selection |
| --- | --- | --- | --- |
| single-copy | 3459 | 539 | 161 |
| ancestral | 19 | 1 | 0 |
| derived | 18 | 2 | 1 |
| relocated | 28 | 4 | 1 |

Relaxed constraints on duplicated genes may be present prior to duplication and not necessary be a result of duplication (O’Toole *et al.*, 2018). To test for relaxed constraints prior to duplication/relocation, we examined the evolution of the *D. melanogaster* orthologs of *D. pseudoobscura* duplicated and relocated genes. We find that *d*_*N*_ /*d*_*S*_ of the *D. melanogaster* orthologs of *D. pseudoobscura* duplicated genes is significantly higher than *d*_*N*_ /*d*_*S*_ of single-copy non-relocated genes, and *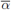* of the orthologs of duplicated genes is greater than the 95% CI of 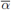of the single-copy genes (**Supplementary Fig S1**). This is consistent with the hypothesis that the ancestors of duplicated genes are more likely to be evolving fast because of positive selection than the ancestors of single-copy genes. In contrast, primate duplicated genes are more likely to have single-copy orthologs that evolve under relaxed constraints, not positive selection (O’Toole *et al.*, 2018). The *D. melanogaster* orthologs of *D. pseudoobscura* relocated genes also have elevated *d*_*N*_ /*d*_*S*_ relative to single-copy non-relocated genes, but 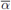 of the orthologs of relocated genes is not higher than 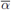 of non-relocated genes (**Supplementary Fig S1**). We therefore conclude that genes evolving under relaxed constraints are more likely to be relocated, but genes that experience positive selection are more likely to be duplicated in *Drosophila*.

### Relocated genes are broadly expressed, are highly expressed in testis, and have male-biased expression

The derived copies of *D. melanogaster* inter-chromosome-arm duplicated genes tend to be narrowly expressed in male reproductive tissues, whereas relocated genes are broadly expressed (Meisel *et al.*, 2009). Using available microarray data from 15 adult *D. melanogaster* tissues, we confirmed that the derived copies of *D. melanogaster* duplicated genes in our data set are more narrowly expressed (higher *τ*) than single-copy genes (**Fig 3A**). In contrast, relocated genes do not significantly differ in their expression breadth from single-copy non-relocated genes (**Fig 3A**) or the ancestral copies of duplicated genes (*P* =0.900 in a Mann-Whitney *U* test).

The derived copies of duplicated genes in animal genomes are often testis-expressed (Vinckenbosch *et al.*, 2006; Meisel *et al.*, 2009, 2010; Baker *et al.*, 2012). We indeed find that the derived copies of *D. melanogaster* duplicates in our dataset are more highly expressed in testis than single-copy non-relocated genes (**Fig 3B**) and the ancestral copies of duplicated genes (*P* =2.8*×*10^−3^ in a Mann-Whitney *U* test). In addition, *D. melanogaster* relocated genes are also more highly expressed in testis than non-relocated genes (**Fig 3B**). Suprisingly, the derived copies of *D. pseudoobscura* duplicated genes are not more highly expressed in testis than either the ancestral copies or single-copy genes (**Fig 3C**). *D. pseudoobscura* relocated genes, on the other hand, are more highly expressed in testis than non-relocated single-copy genes (**Fig 3C**). We therefore conclude that *Drosophila* relocated genes are highly expressed in testis, but the testis-expression of the derived copies of duplicated genes is species-dependent.

**Figure 3:**
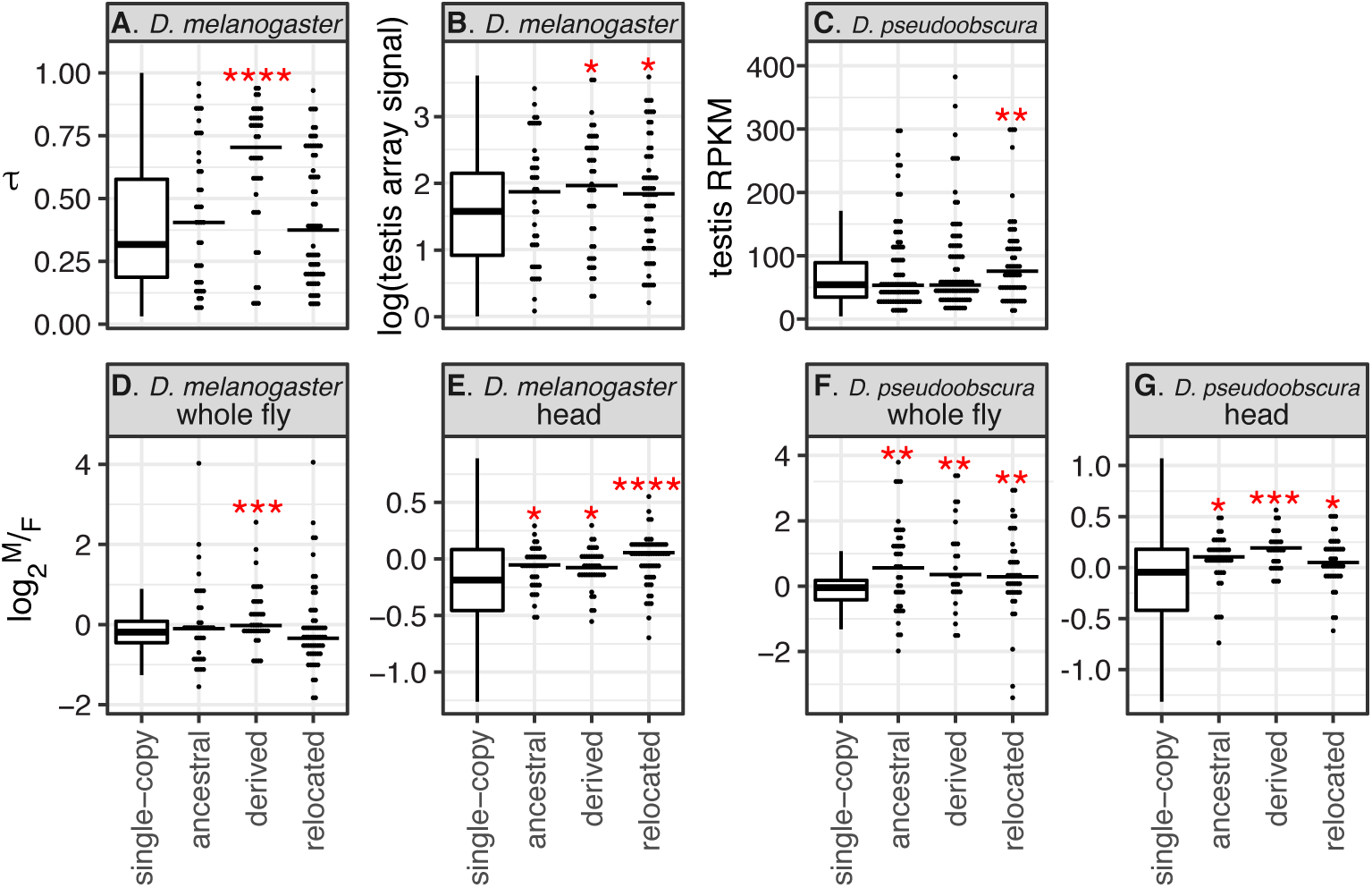
Expression of single-copy non-relocated genes, the ancestral and derived copies of interchromosome-arm duplicated genes, and relocated genes are plotted. (A) Distributions of *τ* for *D. melanogaster* genes are plotted. Distributions of gene expression in testis from (B) *D. melanogaster* microarray data and (C) *D. pseudoobscura* RNA-seq data are plotted. Distributions of log_2_ 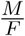 in (D) *D. melanogaster* whole fly, (E) *D. melanogaster* head, (F) *D. pseudoobscura* whole fly, and (G) *D. pseudoobscura* head are plotted. The distribution of log_2_ 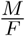 for single-copy genes is represented by a boxplot, while individual values are shown for each of the other genes as a point (with the median indicated by a horizontal line). Significant differences in *τ* when comparing single-copy genes with either ancestral copies, derived copies, or relocated genes are shown by red asterisks (\**P >* 0.05, ^**^*P >* 0.005, ^***^*P >* 0.0005, and ^****^*P >* 0.00005 in a Mann-Whitney *U* test).

Testis expression is the primary driver of male-biased gene expression in *Drosophila* (Parisi *et al.*, 2003). In addition, male-biased and testis expression are among the best predictors of evolutionary rates of protein coding genes (Meisel, 2011). Because relocated genes evolve fast (**Fig 2A–C**) and are testis expressed (**Fig 3B–C**), we assessed whether relocated genes also have male-biased expression. *D. melanogaster* relocated genes do indeed have more male-biased expression than single-copy non-relocated genes in head, but not in whole fly (**Fig 3D–E**). *D. pseudoobscura* relocated genes also have more male-biased expression than single-copy non-relocated genes in head, and in whole fly as well (**Fig 3F–G**). *Drosophila* relocated genes therefore are broadly expressed across many tissues, are highly expressed in male-limited tissues, and have elevated expression in males (**Fig 3**). However, unlike the derived copies of duplicated genes, relocated genes do not have limited expression in male-specific tissues.

To assess how the expression profiles of relocated genes affect their rates of evolution, we calculated Spearman’s non-parametric rank order correlation (?) between each of our divergence estimates and expression metrics for *D. melanogaster* single-copy genes, ancestra opies of inter-chromosome-arm duplicates, derived copies, and relocated genes. Consistent with previous results (Meisel, 2011), faster evolution of single-copy non-relocated genes is associated with more male-biased expression in whole fly (higher log_2_ 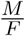), narrower expression (greater *τ*), and higher testis expression (**Supplementary Fig S2**). Faster evolution of relocated genes is also positively correlated with narrower expression (**Supplementary Fig S2**), even though relocated genes are not narrowly expressed (**Fig 3A**). In contrast, testis expression levels are not positively correlated with evolutionary rate for relocated genes (**Supplementary Fig S2**), even though relocated genes evolve fast (**Fig 2A-C**) and are highly expressed in testis (**Fig 3B**). We observe similar results for the derived copies of inter-chromosome-arm duplicated genes (**Supplementary Fig S2**). We therefore conclude that higher testis expression could explain the faster evolution of relocated genes when compared to single-copy non-relocated genes, but expression breadth is the best predictor of evolutionary rates within relocated genes.

### Relocated genes are not disproportionately essential for viability

The broad expression of relocated genes suggests that they may be essential for viability. To test this hypothesis, we compared the effects of RNAi knockdown of relocated genes to knockdown of single-copy non-relocated genes in *D. melanogaster*. We first analyzed the effect of knockdown using randomly inserted UAS-RNAi constructs (GD lines) that do not have any known systemic effect on viability independent of RNAi knockdown of the target gene. Using those data, we find some evidence that relocated genes are less essential than non-relocated single-copy genes. Knockdown of less than a quarter (5*/*24) of relocated genes causes a significant decrease in viability (**Fig 4A**), whereas over half (8*/*15) of the single-copy non-relocated genes have a significant viability effect when knocked down (**Fig 4B**; *P* =0.079 in Fisher’s exact test). In addition, we quantified the effect of knockdown on viability, with more negative values indicating a larger effect. The median effect of knockdown on viability is significantly more negative for single-copy non-relocated genes than relocated genes (*P* =1.810^−4^ in a Mann-Whitney *U* test).

We also tested the effect of knockdown using site-specific UAS-RNAi construct insertions (KK lines) that have a known systemic effect on viability (Green *et al.*, 2014; Vissers *et al.*, 2016). Indeed, we observe that the median knockdown effect on viability is more negative using the KK lines (*-*13.54; see **Supplementary Fig S3**) than the GD lines (1.13; **Fig 4**) for relocate genes (*P* =0.005 in a Mann-Whitney *U* test). Therefore, the systemic effects of the KK lines on viability make them unsuited for inferring the essentiality of both relocated and duplicated genes (Kondo *et al.*, 2017). Surprisingly, there is not a more negative viability effect of knockdown using KK lines (*-*15.53) than GD lines (*-*20.36) for single-copy non-relocated genes.

The broader expression breadth of relocated genes than duplicated genes (**Fig 3A**) suggests that relocated genes are more likely to be essential for viability. To test this hypothesis, we compared our results examining the effect of knockdown of relocated and non-relocated genes with published data assessing if ubiquitous knockdown of the derived copies of *D. melanogaster* duplicated genes induces lethality (Chen *et al.*, 2010). In this approach, if knockdown of a gene causes lethality, the gene is considered essential. We considered knockdown of relocated and non-relocated genes to induce lethality if there were no knockdown progeny recovered in at least 90% of replicate experiments we performed with at least one GD RNAi line, similar to the criteria for considering the lethal effect of knocking down duplicated genes in the published data (Chen *et al.*, 2010). We only considered inter-chromosome-arm duplications, and we only analyzed results from GD lines because of the systemic effects of KK lines described above and previously reported (Green *et al.*, 2014; Vissers *et al.*, 2016; Kondo *et al.*, 2017). We observe that similar proportions of relocated genes (16.7%) and derived copies of duplicated genes (15.0%) cause lethality when knocked down (**Table 2**). In contrast, 40% of single-copy non-relocated genes causes lethality (**Table 2**), but this is not significantly different from the fraction of essential relocated genes (*P* =0.14 in Fisher’s exact test) or duplicated genes (*P* =0.13 in Fisher’s exact test). We therefore conclude that, despite the increased expression breadth of relocated genes, they are not more likely to be essential for viability than the derived copies of duplicated genes. We also observe that, for both relocated and non-relocated genes, there is not a significant correlation between expression breadth and the effect of knockdown on viability (**Supplementary Fig S4**). These results suggest that expression breadth is not a reliable proxy for gene essentiality.

**Figure 4:**
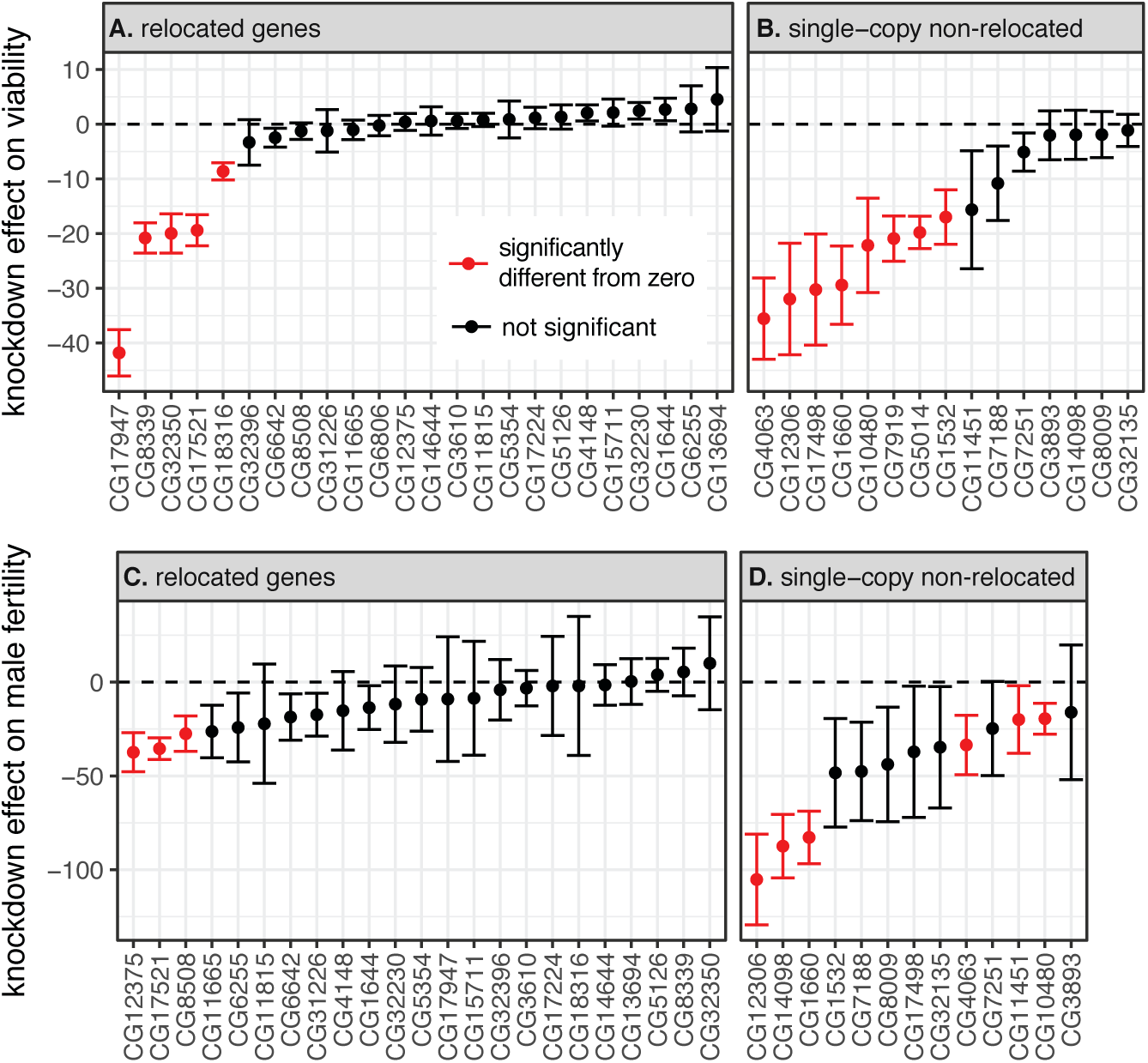
The effects of RNAi knockdown on viability and fertility are plotted. Knockdown was performed using (A–B) ubiquitous expression of Gal4 to assess viability and (C–D) germline expression of Gal4 to assess male fertility. Only data using GD lines are plotted. RNAi targeted (A & C) relocated genes or (B & D) single-copy non-relocated genes. Dots indicate the mean effect of knockdown across replicates, and the vertical bars show the standard error. Each point is a gene, and those colored red have knockdown effects significantly less than zero, indicating decreased in (A–B) viability or (C–D) male fertility.

**Table 2:** Counts of relocated genes, derived copies of inter-chromosome-arm duplicates, and singlecopy non-relocated genes that are lethal or non-lethal to knockdown. The criterion for lethality is no knockdown progeny recovered in 90% of replicate experiments.

|  | lethal | non-lethal | perc lethal |
| --- | --- | --- | --- |
| Relocated | 4 | 20 | 16.7% |
| Derived dups | 3 | 17 | 15.0% |
| Single-copy | 6 | 9 | 40.0% |

The genes included in the viability assays are a subset of all single-copy, duplicated, and relocated genes in the *D. melanogaster* genome. We tested if they are representative of the patterns of divergence, selection, and expression we observe in the full set of single-copy, duplicated, and relocated genes. We confirmed that the derived copies of duplicated genes that were included in the viability assays do indeed evolve faster than the single-copy non-relocated genes included in the viability assays at all classes of sites (**Supplementary Fig S5**). The relocated genes included in our viability assays also evolve faster than the single-copy non-relocated genes at 4-fold degenerate sites (**Supplementary Fig S5**). In addition, the derived copies of duplicated genes in our viability assays are narrowly expressed, and they have more male-biased expression (greater log_2_ 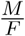) in whole fly than the single-copy genes (**Supplementary Fig S5**). The relocated genes included in the viability assays also have more male-biased expression than the single-copy non-relocated genes (**Supplementary Fig S5**). We therefore conclude that the genes included in the viability assays are in general representative of single-copy, duplicated, and relocated genes in the *D. melanogaster* genome.

### Relocated genes are not disproportionately essential for male fertility

Relocated genes are highly expressed in testis (**Fig 3B–C**), suggesting that their products may perform essential roles in spermatogenesis. To test this hypothesis, we assessed the fertility of *D. melanogaster* males in which relocated genes were knocked down in the male germline using GD UAS-RNAi lines. We compared our results to germline knockdown of single-copy non-relocated genes. We also quantified the effect of knockdown on male fertility, with more negative values indicating a larger decrease in male fertility relative to controls.

Surprisingly, despite their higher testis expression, we do not find evidence that relocated genes are more essential for male fertility than single-copy non-relocated genes. Germline knockdown of only 3*/*23 relocated genes induced a significant decrease in male fertility (**Fig 4C**), compared to nearly half (6*/*13) of single-copy non-relocated genes (**Fig 4D**; *P* =0.046 in Fisher’s exact test). In addition, the median knockdown effect on male fertility is more negative for non-relocated genes than relocated genes (**Fig 4C–D**; *P* =1.8*×*10^−5^ in a Mann-Whitney *U* test).

We also tested the effect of knockdown on male fertility using site-specific UAS-RNAi construct insertions (KK lines) that have a known systemic effect on viability (Green *et al.*, 2014; Vissers *et al.*, 2016). Unlike the viability effects, we do not detect a systemic effect of the KK lines on male fertility (**Supplementary Fig S3**). In fact, the effect of knockdown on male fertility using GD lines (*-*9.21) is more negative than with KK lines (6.52) for relocated genes (*P<*0.01 in both paired and unpaired Mann-Whitney *U* tests). The knockdown effect on male fertility for single copy genes is also more negative for GD lines (*-*40.5) than KK lines (*-*8.58; *P* =0.039 in a paired Mann-Whitney *U* test). Our results therefore suggest that both GD and KK lines can be used to assess if genes are necessary for male fertility. In addition, male fertility under germline knockdown is negatively correlated with testis expression level for both relocated and non-relocated genes (**Supplementary Fig S6**), suggesting that testis expression is predictive of the effects of germline knockdown.

Finally, we tested if the genes included in the fertility assay are a representative subset of all non-relocated and relocated genes in the *D. melanogaster* genome. The relocated genes in our fertility assay evolve faster at 4-fold degenerate sites than the non-relocated genes, and they have more male-biased expression than the non-relocated genes (**Supplementary Fig S7**). However, the relocated genes in our fertility assay are not more highly expressed in testis than the non-relocated genes (**Supplementary Fig S7**). This is because both the relocated and non-relocated genes included in our fertility assay have higher testis expression than the genes not included in the fertility assay (*P* =0.024 and *P* =0.0019 in a Mann-Whitney *U* test for relocated and non-relocated genes, respectively). It is therefore possible that the result of our fertility assay was biased by selecting relocated and non-relocated genes with higher testis expression than average.

## Discussion

Gene relocation occurs frequently in eukaryotic genomes, and relocated genes may be as common as canonical inter-chromosomal duplicated genes (Bhutkar *et al.*, 2007; Wicker *et al.*, 2010; Han and Hahn, 2012). In addition, relocated genes have been hypothesized to evolve under more selective constraints than the derived copies of dispersed duplicated genes, and they may be as conserved as single-copy non-relocated genes (Meisel *et al.*, 2009; Ciomborowska *et al.*, 2013). We found that relocated genes evolve faster than non-relocated genes, and there is no evidence that this faster evolution is driven by positive selection (**Fig 2** & **Table 1**). We therefore conclude that relocated genes evolve fast because they are under relaxed constraints. The derived copies of dispersed duplicates evolve even faster than relocated genes, consistent with the hypothesis that relocated genes are under more selective constraints than duplicated genes.

Relocated genes are broadly expressed, while the derived copies of inter-chromosome-arm duplicates are narrowly expressed (**Fig 3**). Broad expression and high expression levels are associated with slower evolution of *Drosophila* genes (Larracuente *et al.*, 2008; Meisel, 2011). It is therefore not surprising that the derived copies of inter-chromosome-arm duplicates evolve faster than both non-relocated and relocated genes (**Fig 2**). However, despite their broad expression, relocated genes also evolve faster than non-relocated genes (**Fig 2**). Our results suggest that, even though expression level and breadth are predictive of evolutionary rates within categories of genes (Larracuente *et al.*, 2008; Meisel, 2011, **Supplementary Fig S2**), expression differences between categories of genes are poorly associated with evolutionary rate differences between categories.

The broad expression and high testis expression of relocated genes led us to hypothesize that they are essential for viability and male fertility (**Fig 3**). However, our RNAi experiments revealed that the relocated genes are less essential for viability and male fertility than single-copy non-relocated genes (**Fig 4** & **Table 2**). This is consistent with our results that suggest relocated genes evolve faster than single-copy non-relocated genes because relocated genes are under relaxed selective constraints (**Fig 2** & **Table 1**). These results also demonstrate that functional analyses that complement expression measurements are necessary to identify differences in selective constraints acting on different classes of genes.

Our inference of relaxed constraints comes, in part, from analyses of extant DNA sequences and effects of RNAi knockdown on extant relocated genes. It is possible that relocated genes (and duplicated genes) could have experienced strong positive or purifying selection immediately after duplication (or loss of the ancestral paralog), and the signatures of those selection pressures were lost over time. Analysis of a large panel of young duplications and relocations are necessary to test this hypothesis (e.g., Masly *et al.*, 2006; VanKuren and Long, 2018). However, our observation that the *D. melanogaster* orthologs of *D. pseudoobscura* relocated genes also evolve under relaxed constraints (**Supplementary Fig S1**) suggests that relocated genes have evolved under relaxed constraints for most of their histories.

An excess of genes has been relocated from the X chromosome to the autosomes across the *Drosophila* genus (Meisel *et al.*, 2009; Vibranovski *et al.*, 2009b). Three hypotheses could explain this phenomenon. First, the female-biased transmission of the X chromosome may favor X-linked female-beneficial mutations and prevent the fixation of male-beneficial mutations on the X (Rice, 1984). This sexually antagonistic selection could favor the X-to-autosome relocation of genes that perform male-beneficial functions (Wu and Xu, 2003). Second, expression of the X chromosome is down-regulated in spermatogenesis (Vibranovski *et al.*, 2009a; Meiklejohn *et al.*, 2011), which could favor the X-to-autosome relocation of genes that have beneficial effects when highly expressed in spermatogenesis (Betrán *et al.*, 2002; Emerson *et al.*, 2004; Meisel *et al.*, 2009). Third, there may be a mutational bias in favor of X-to-autosome duplications (Metta and Schlotterer, 2010; Díaz-Castillo and Ranz, 2012), but this is not supported by copy number polymorphisms (Schrider *et al.*, 2011). We find no evidence that knockdown of relocated genes disproportionately affects male fertility (**Fig 4**). However, relocated genes are more highly expressed in testis than non-relocated single-copy genes (**Fig 3**), which holds true even if we only consider autosomal genes (*P <* 0.05 for both *D. melanogaster* and *D. pseudoobscura*). Our gene expression analysis therefore provides some evidence that the X-to-autosome relocation bias could be driven by selection in favor of higher testis expression on the autosomes, but additional work is necessary to fully test this hypothesis.

There are two important technical limitations of our experiments that reduce our ability to detect male-specific functions of relocated genes. First, we used ubiquitous and germline knockdown to assess if genes are essential for viability and male fertility. We chose to assay the effect of germline knockdown because relocated genes are highly expressed in testis (**Fig 3**), and the derived copies of duplicated genes are hypothesized to be specialized for germline functions (Marques *et al.*, 2005; Vinckenbosch *et al.*, 2006; Potrzebowski *et al.*, 2008; Meisel *et al.*, 2010; Tracy *et al.*, 2010). We demonstrated that, even though they are highly expressed in testis, relocated genes are not disproportionately essential for spermatogenesis (**Fig 4**). However, our results may be biased by the Gal4 driver that we selected (*bam*), which is expressed early in spermatogenesis immediately after differentiation from the stem cell niche (Chen and McKearin, 2003). Knockdown in later stages of spermatogenesis or in somatic testis tissue may reveal testis-biased functions for relocated genes. Second, germline knockdown of a panel of duplicated genes would allow for a comparison of male-specific functions between relocated genes and derived copies of inter-chromosome-arm duplicates. This may reveal additional insights into the causes of differences in the evolutionary rates of duplicated and relocated genes (**Fig 2**).

In conclusion, we demonstrated that *Drosophila* relocated genes evolve fast, and this rapid evolution is likely the result of relaxed selective constraints (**Fig 2** & **Table 1**). This differs from mammals, where relocated genes evolve under strong purifying selection (Ciomborowska *et al.*, 2013). *Drosophila* relocated genes are also less essential for viability and male fertility than single-copy non-relocated genes (**Fig 4**), which is consistent with relocated genes evolving under relaxed constraints. In addition, the derived copies of inter-chromosome-arm duplicates appear to be under even more relaxed constraints than relocated genes, which allows them to evolve even faster. Additional work is necessary to determine the causes of differences in selection pressures acting on relocated genes in mammals and *Drosophila*.

## Acknowledgments

We thank members of the Meisel lab at the University of Houston and Andy Clark’s lab at Cornell University for assistance with the RNAi experiments. Mariana Wolfner kindly supplied the *bam-Gal4* line, which was originally produced in Margaret Fuller’s laboratory. Erin Kelleher provided valuable feedback on the preparation of this manuscript. This work was supported by start up funds from the University of Houston to RPM and a University of Houston Summer Undergraduate Research Fellowship to LZ.

## Supplemental Figures and Tables

**Supplemental Fig S1:**
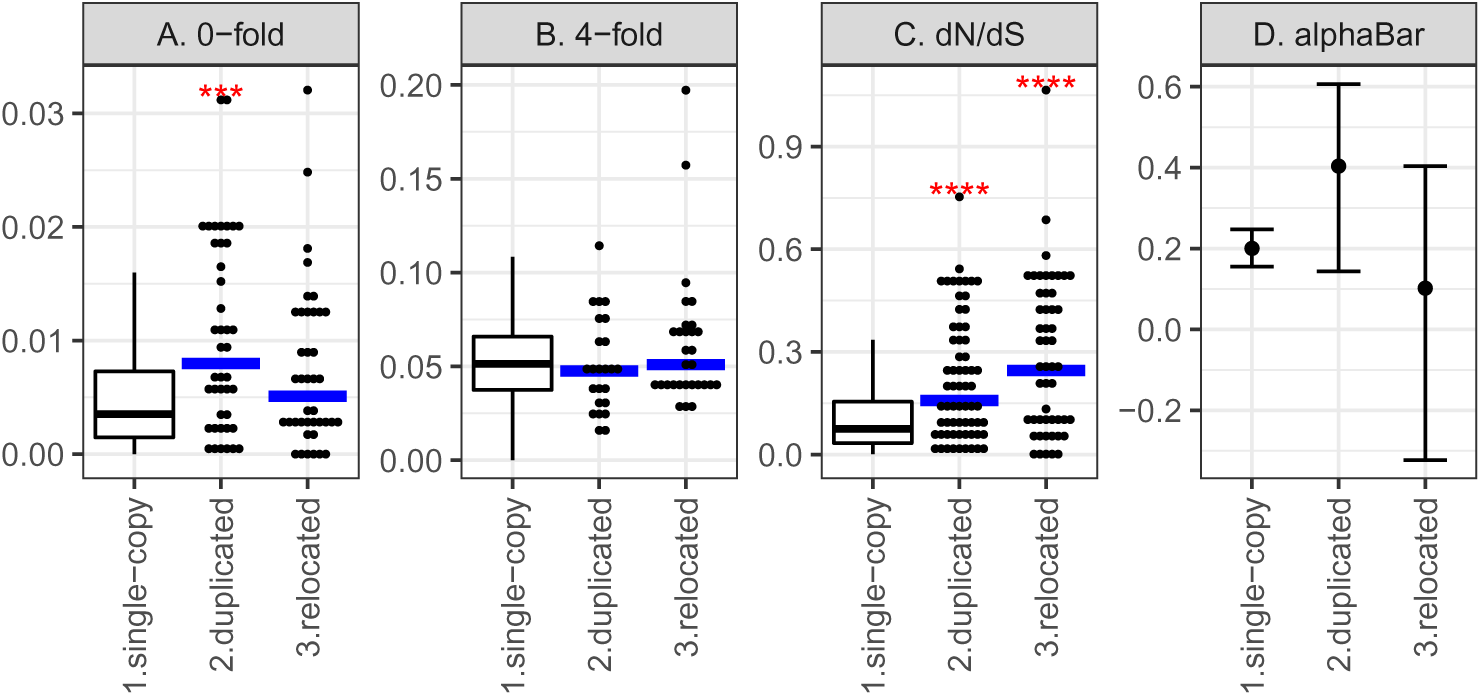
Divergence measures and tests for selection are plotted for *D. melanogaster* homologs of *D. pseudoobscura* single-copy, duplicated, and relocated genes. For all metrics except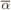, the distribution of divergence values for single-copy genes is represented by a boxplot, and individual divergence values are shown for each of the other genes as a point (with the median indicated by a horizontal blue line). Estimates of 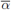are plotted as a point, along with the 95% CI. Significant differences in values when comparing single-copy non-relocated genes with duplicated and relocated genes are shown by red asterisks (^***^*P >* 0.0005, and ^****^*P >* 0.00005 in a Mann-Whitney *U* test).

**Supplemental Fig S2:**
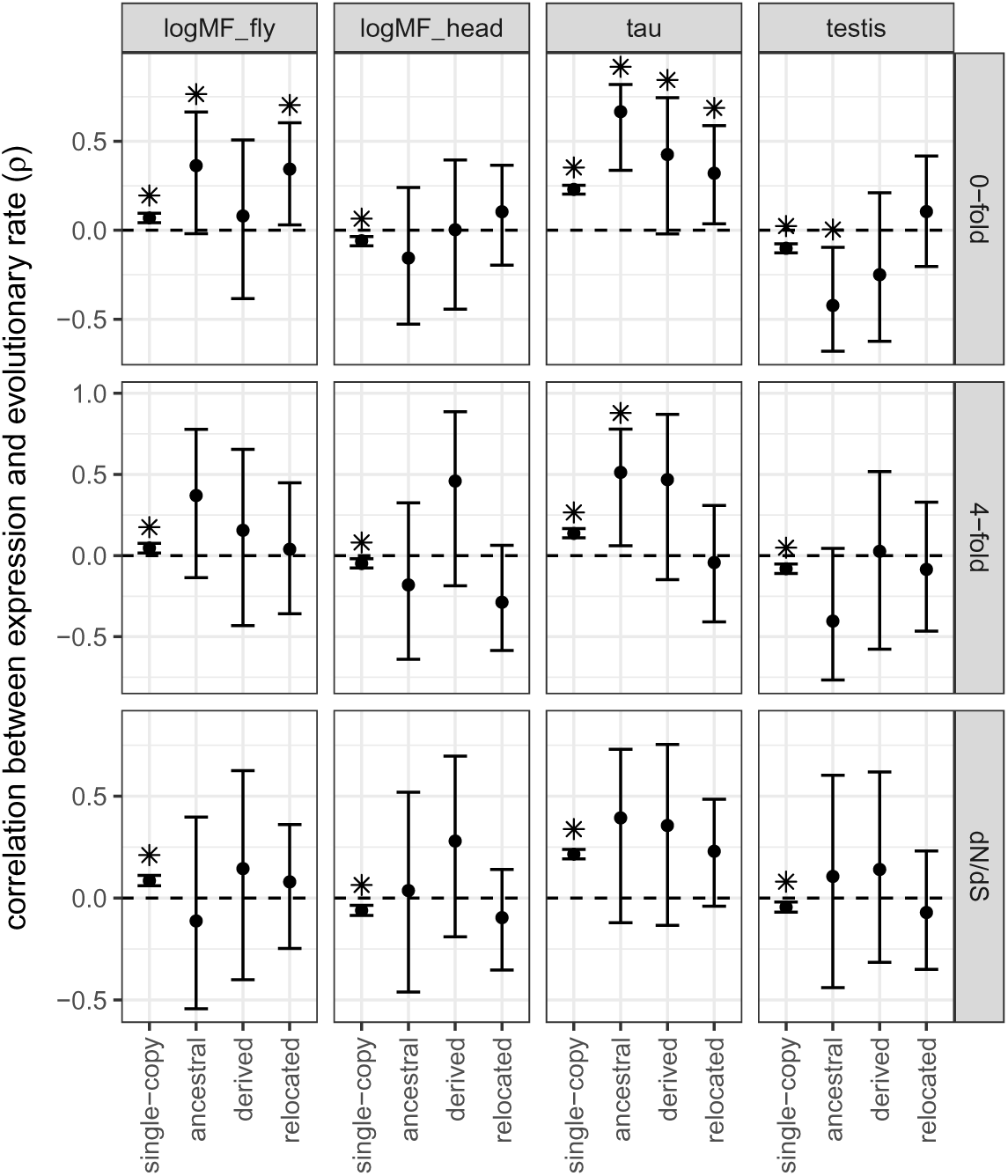
Spearman’s non-parametric rank order correlation (*?*) between all pair-wise combinations of expression metrics (columns) and evolutionary divergence (rows) is plotted for single-copy non-relocated genes, the ancestral and derived copies of inter-chromosome-arm duplicates, and relocated genes. Error bars are 95% CIs from 1,000 bootstrap replicates of the data. Asterisks indicate correlations that are significantly different from zero.

**Supplemental Fig S3:**
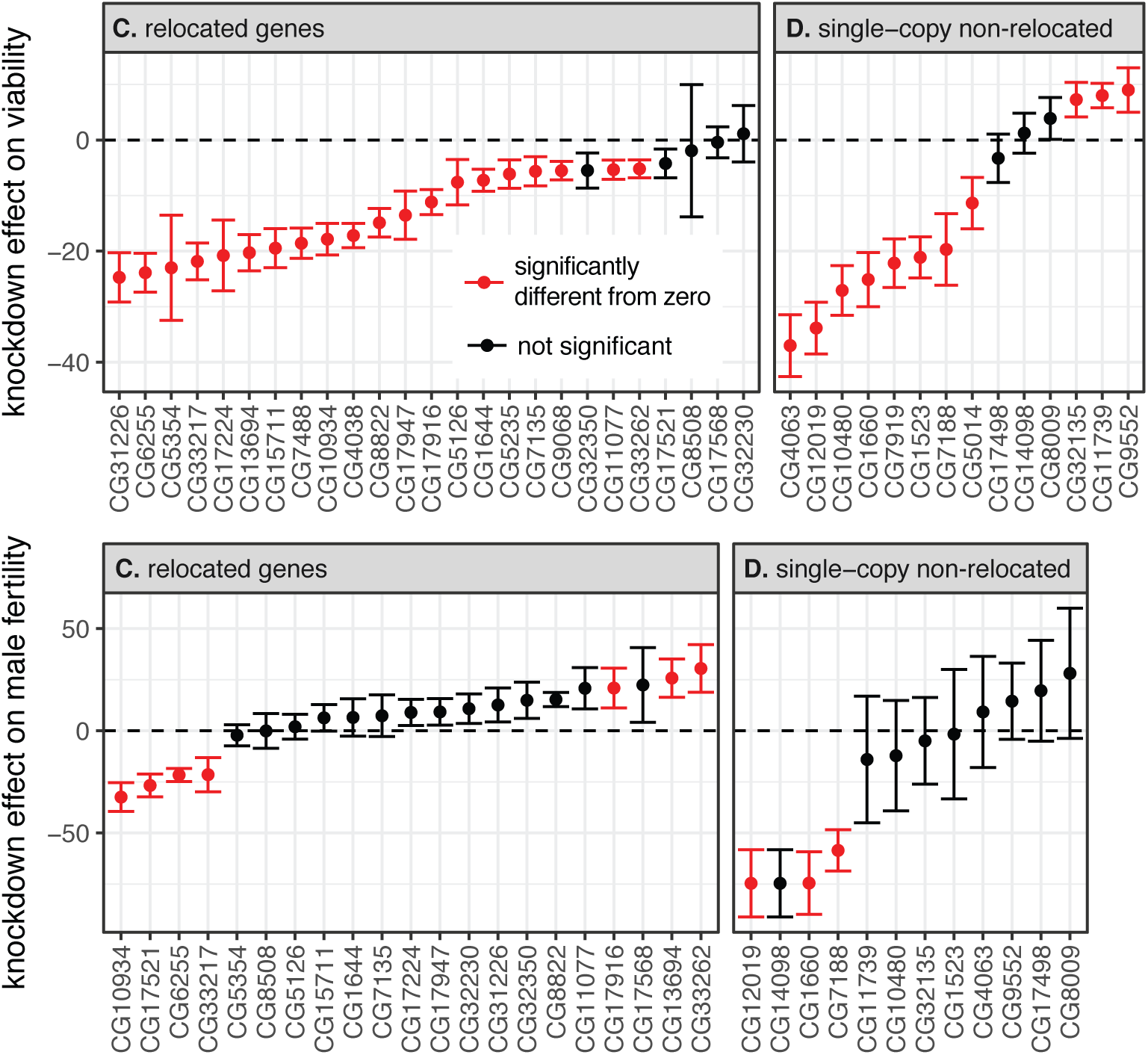
The effects of RNAi knockdown using KK lines on viability and fertility are plotted. Knockdown was performed using (A–B) ubiquitous expression of Gal4 to assess viability and (C–D) germline expression of Gal4 to assess male fertility. Only data using KK lines are plotted. RNAi targeted (A & C) relocated genes or (B & D) single-copy non-relocated genes. Dots indicate the mean effect of knockdown across replicates, and the vertical bars show the standard error. Each point is a gene, and those colored red have knockdown effects significantly less than zero, indicating decreased in (A–B) viability or (C–D) male fertility.

**Supplemental Fig S4:**
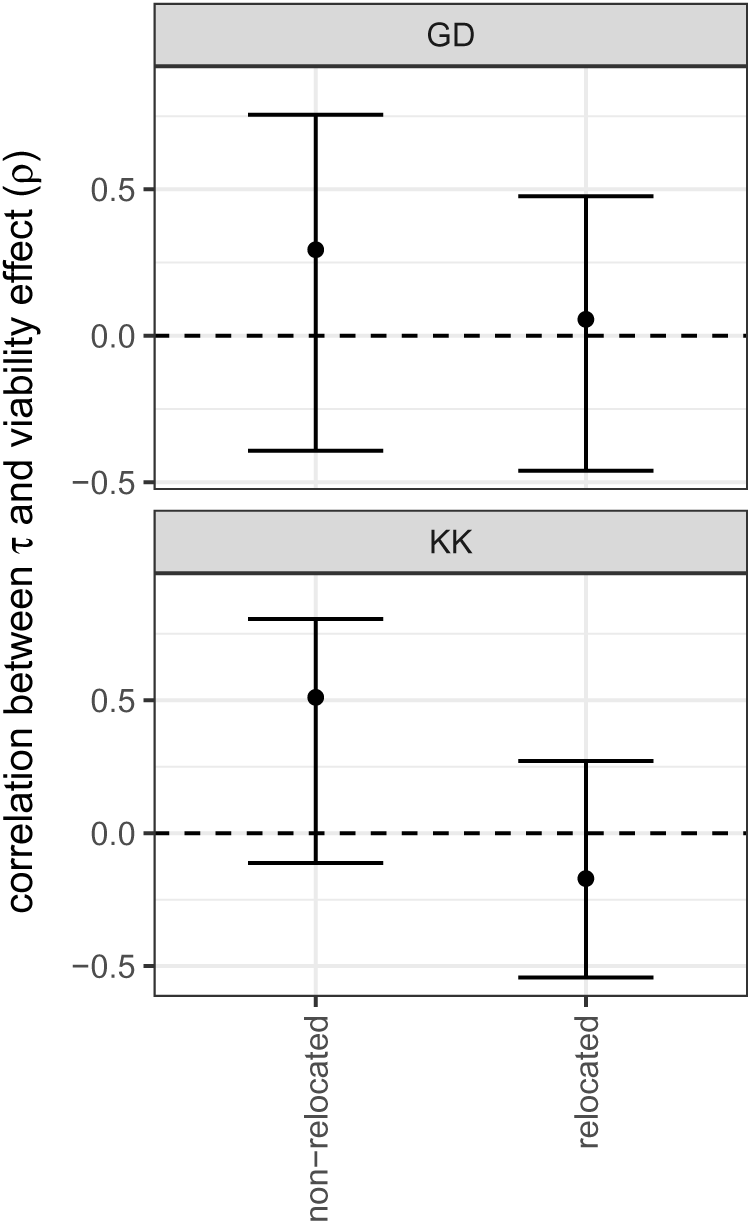
Spearman’s non-parametric rank order correlation (*ρ*) between expression breadth (*τ*) and effect of RNAi knockdown on viability for single-copy non-relocated genes and relocated genes. Viability effects were determined using data from GD lines (top) or KK lines (bottom). Error bars are 95% CIs from 1,000 bootstrap replicates of the data. No correlations are significantly different from zero.

**Supplemental Fig S5:**
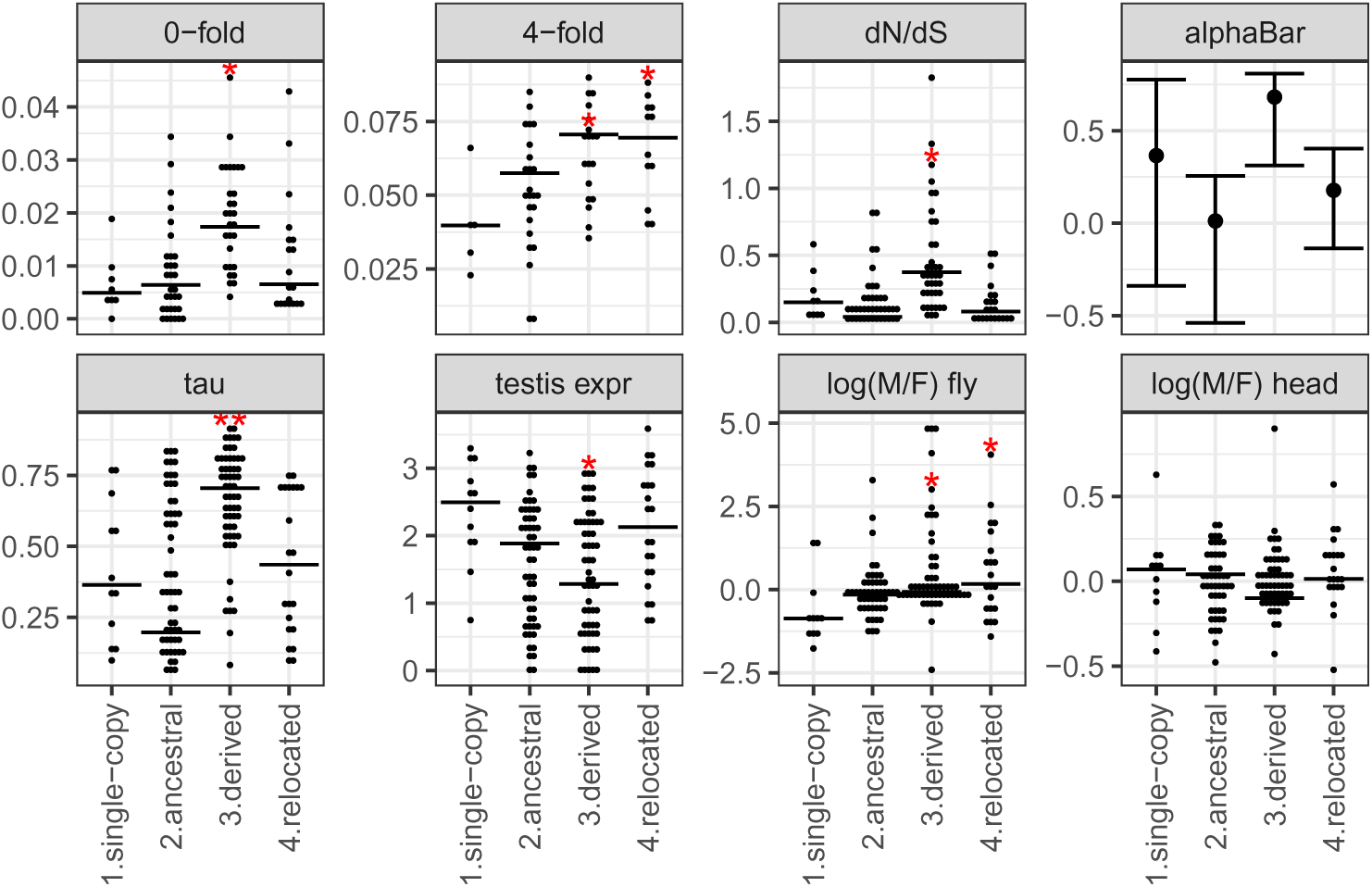
Divergence measures, tests for selection, and gene expression data are plotted for genes used in the RNAi assays of viability effects. Estimates of 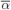are plotted as a point, along with the 95% CI. Significant differences in values when comparing single-copy non-relocated genes with genes in the other three classes are shown by red asterisks (^*^*P >* 0.05, ^**^*P >* 0.005, ^***^*P >* 0.0005, and ^****^*P >* 0.00005 in a Mann-Whitney *U* test).

**Supplemental Fig S6:**
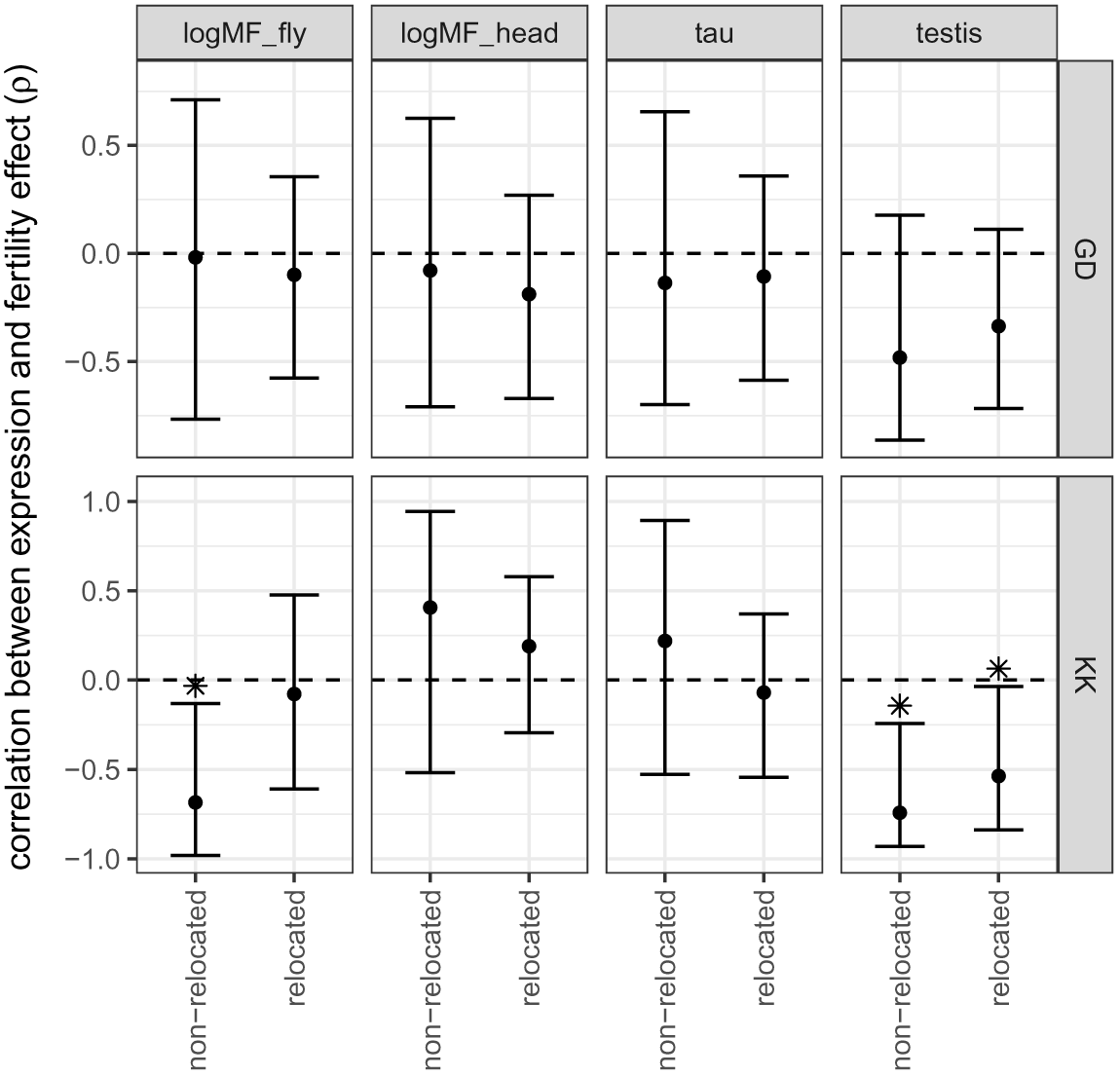
Spearman’s non-parametric rank order correlation (*ρ*) between gene expression measures (columns) and effect of RNAi knockdown on fertility for single-copy non-relocated genes and relocated genes. Fertility effects were determined using data from GD lines (top) or KK lines (bottom). Gene expression is either log_2_ 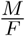 for whole flies, log_2_ 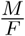 for heads,*τ*, or testis expression level. Error bars are 95% CIs from 1,000 bootstrap replicates of the data. Asterisks indicate correlations that are significantly different from zero.

**Supplemental Fig S7:**
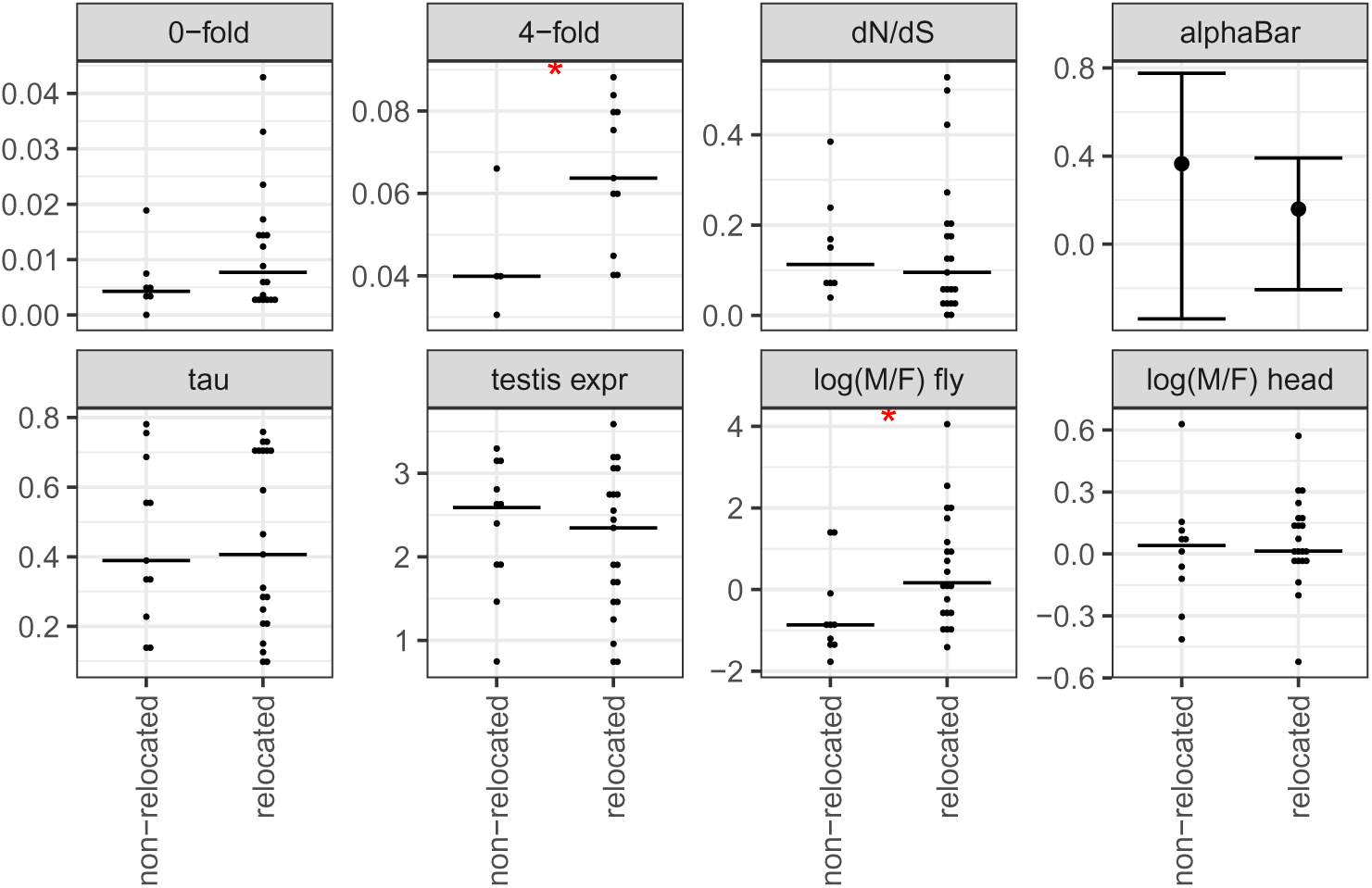
Divergence measures, tests for selection, and gene expression data are plotted for genes used in the RNAi assays of fertility effects. For all metrics except 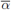each dot is an individual gene, and the median across genes is indicated by a horizontal line. Estimates of 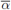are plotted as a point, along with the 95% CI. Significant differences in values when comparing non-relocated genes with relocated genes are shown by red asterisks (^*^*P >* 0.05, ^**^*P >* 0.005, ^***^*P >* 0.0005, and ^****^*P >* 0.00005 in a Mann-Whitney *U* test).

